# Evaluating instruments for assessing healthspan: a multi-center cross-sectional study on health-related quality of life (HRQL) and frailty in the companion dog

**DOI:** 10.1101/2022.07.21.500746

**Authors:** Frances L. Chen, Tarini V. Ullal, Jessica L. Graves, Ellen R. Ratcliff, Alexander Naka, Brennen McKenzie, Tennery A. Carttar, Kaitlyn M. Super, Jessica Austriaco, Sunny Y. Weber, Julie Vaughn, Michael L. LaCroix-Fralish

**Author notes:** **Statements and Declarations** Competing Interests: All authors are employed by Loyal, a part of Cellular Longevity, Inc.

## Abstract

Developing valid tools that assess key determinants of canine healthspan such as frailty and health-related quality of life (HRQL) is essential to characterizing and understanding aging in dogs. Additionally, because the companion dog is an excellent translational model for humans, such tools can be applied to evaluate gerotherapeutics and investigate mechanisms underlying longevity in both dogs and humans. In this multi-center, cross sectional study, we investigated the use of a clinical questionnaire (Canine Frailty Index; CFI; Banzato et al., 2019) to assess frailty and an owner assessment tool (VetMetrica HRQL) to evaluate HRQL in 451 adult companion dogs. Results demonstrated validity of the tools by confirming expectations that frailty and HRQL deteriorate with age. CFI scores were significantly higher (higher frailty) and HRQL scores significantly lower (worse HRQL) in old dogs (≥ 7 years of age) compared to young dogs (≥ 2 and < 6 years of age). Body size (small < 25lbs or large > 50lbs) was not associated with CFI or total HRQL score. However, older, larger dogs showed faster age-related decline in HRQL scores specific to owner-reported activity and comfort. Findings suggest that the clinician-assessed CFI and owner-reported VetMetrica HRQL are useful tools to evaluate two determinants of healthspan in dogs: the accumulation of frailty and the progressive decline in quality of life. Establishing validated tools that operationalize the assessment of canine healthspan is critical for the advancement of geroscience and the development of gerotherapeutics that benefit both human and veterinary medicine.

**Graphical abstract:** Graphical summary of the design, results, and conclusions of the study.

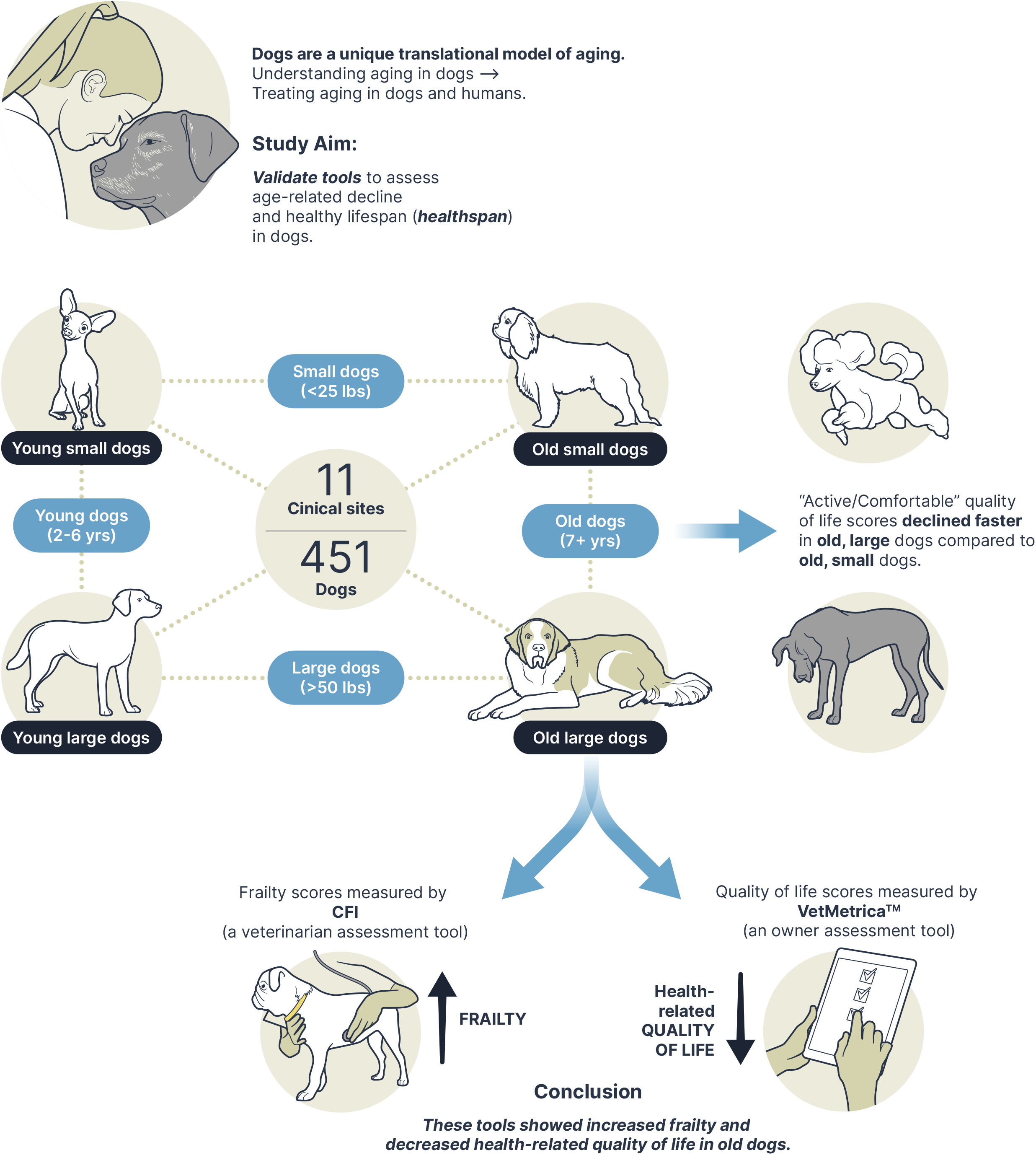

## Introduction

Aging is the single greatest risk factor for nearly every major cause of morbidity and mortality in living organisms [1–4], including companion dogs [5]. Even in the absence of clinical disease, loss of function at the cellular and tissue levels underlies progressive impairments in physiological and homeostatic function of organ systems throughout life [6] that result in declining daily function and performance and increasing multimorbidity. Several studies in model organisms have demonstrated that the aging process is malleable and can be slowed with medical interventions and lifestyle alterations to delay mortality and promote a healthy lifespan or “healthspan” [7–14]. However, in contrast to traditional laboratory models, the domestic dog offers a novel translational opportunity to investigate aging biology and potential gerotherapeutics. Companion dogs live in shared environments with humans and have similar age-associated diseases and access to advanced healthcare. Simultaneously, dogs have the greatest intraspecies phenotypic diversity of any mammal, including a 2-2.5 fold variation in lifespan, which enhances the study of risk factors and biological pathways contributing to longevity. Dogs live shorter lives than humans, making evaluation of gerotherapeutics feasible on a meaningful timescale. Most importantly, companion dogs are seen as valued family members and there is a strong societal drive to increase their healthspan and lifespan. For all these reasons, comparative and translational research in the dog holds the promise of identifying novel strategies that mitigate aging and improving health outcomes in both canines and humans [15,16].

Quantitative outcome measures to assess the multidimensional process of aging and multifactorial components of healthy lifespan in the companion dog are necessary for advancing mechanistic understanding of aging biology and selecting clinical trial endpoints for candidate gerotherapeutics. An important approach for measuring the impact of aging is to assess *healthspan*, which we define here as “the period of life spent in good health, free from the chronic diseases and disabilities of aging.” In order to measure healthspan (i.e. quantification of the period of time the organism is free from decline), one must be able to quantify the health-related features that change in an organism due to aging. Two important features that can be measured to quantify healthspan include assessments of *frailty*, a state of vulnerability and diminished physiological reserve, and health-related quality of life (HRQL), the perceived impact of one’s health on overall well-being. Because there is currently no clear consensus on gold standard methods for assessing *healthspan* in dogs (Kaeberlein, 2018), operationalizing tools that do so are critical to capturing the negative impact of aging in dogs and advancing translational geroscience. Gerontology research in humans has led to the development of several tools that quantify components of healthspan-e.g. through patient-reported or clinical outcome measures of frailty, quality of life, physical function/capacity, and activity [17–20]. Only recently has there been an interest to develop analogous outcome measures in veterinary gerontology. Here, we selected two such tools (Canine Frailty Index (CFI) and the VetMetrica Health Related Quality of Life (HRQL) instrument) to investigate their utility in evaluating two important readouts of healthspan in the companion dog: frailty and health-related quality of life.

The Canine Frailty Index (CFI) was developed to measure frailty in dogs by scoring age-associated multimorbidity and functional deficits and determining the relationship of that score to short term mortality risk [21]. The CFI applied in dogs is analogous to the frailty index used in humans or rodents where frailty is represented as a general state of poor health and higher risk of mortality proportional to the number of age-associated deficits accumulated [22,23]. The CFI consists of 33 discrete clinical health parameters from difficulty climbing stairs to oral disease assessed by a veterinarian, which are added and divided by the total number of items to create a composite score. Higher CFI scores indicate a higher number of age-associated diseases, deficits, and overall greater frailty, which is associated with increased short-term (6 month) mortality risk [21]. In a prospective study in 401 dogs, CFI score was moderately correlated with age and the distribution of scores also increased with age [21].

In human clinical trials, patient-reported outcomes for health-related quality of life (HRQL) are critical to a patient-centric approach for assessing benefit of an intervention. Analogously, in veterinary clinical trials, valid owner assessed outcome measures ensure meaningful interpretation when evaluating efficacy of an intervention. These instruments must undergo proper psychometric testing and validation. The VetMetrica^TM^ Health Related Quality of Life (HRQL) instrument was developed by NewMetrica Ltd. to assess general health related quality of life of companion dogs as perceived by the pet owner. The VetMetrica instrument is an online questionnaire that covers four domains of health-related quality of life in companion dogs: Energetic/Enthusiastic (E/E), Happy/Content (H/C), Active/Comfortable (A/C), and Calm/Relaxed (C/R) and has previously been psychometrically validated [24,25]. VetMetrica instantaneously computes and reports a profile of raw scores across the 4 domains [26]. Importantly, VetMetrica HRQL has previously shown lower quality of life scores in older dogs [27] as well as in dogs with illness [24,26].

In addition to potentially serving as outcome measures for clinical trial endpoints, assessment tools for healthspan can be used to identify risk factors and explore hypotheses on biological pathways that contribute to healthspan and lifespan differences. A number of studies have now established a consistent inverse correlation found between median lifespan and adult body mass in dogs [28–30]. However, it is currently unclear why large dogs have shorter lifespans compared to small dogs. A possible explanation is that large and giant breed dogs have higher mortality risk with increasing age [31] compared to small dogs, but not necessarily differences in multimorbidity [21,31,32]. This theoretical model of ‘truncation’ (shortened lifespan without accelerated aging) or ‘compression’ (accelerated aging leading to shortened lifespan) in large sized dogs has been previously discussed and is an ongoing area of investigation [16,33]. Tools that are sensitive enough to detect biologically meaningful differences between subpopulations of dogs (such as small and large) but are still applicable to all dogs could be used as clinical trial endpoints or in observational, investigative research to determine if and how biological pathways underlying body size contribute to aging in dogs.

While both the CFI and the VetMetrica HRQL instruments have undergone initial development and validation, further evaluation and refinement in large scale companion dog populations and multi-center studies are necessary to determine their potential utility in measuring aging-related decline and assessing potential gerotherapeutics in companion dogs. Here we report a cross-sectional, multi-site veterinary clinical study that aimed to assess the utility of the clinically-assessed CFI and the owner-reported VetMetrica HRQL instrument in detecting differences in health-related quality of life and frailty between young (≥ 2 and < 6 years of age) and old (≥ 7 years of age) dog groups. Specifically, we hypothesized that the CFI and VetMetrica HRQL tools would show increased frailty and decreased HRQL scores in older dogs. We also hypothesized that owner-assessed HRQL scores would show an inverse relationship with clinician-assessed CFI scores, demonstrating that owner-assessed HRQL relates to the clinical status of the dog as assessed by a veterinarian (CFI). Finally, given the evidence that dogs of large body size have shorter expected lifespans [28–30], we also assessed potential differences in CFI and HRQL between old and young dogs of small (<11.3 kg ; <25 lbs) and large (> 22.7 kg ; 50 lbs) body size to evaluate whether larger dogs show increased frailty or worse health-related quality of life with age compared to smaller dogs.

## Materials and Methods

### IACUC approval, Study Standards, and Animal Welfare

The study was performed according to VICH-GL9 Good Clinical Practice (GCP) standards. Approval by an independent IACUC was obtained on 13 Dec 2020 through Veterinary & Biomedical Research Center, Inc (VBRC) (Approval #: VACLOY001CLDEFF1PILC). Owner consent was obtained prior to any study procedures.

### Study Design & Experimental Subjects (Fig. 1)

This study was a multi-center, cross-sectional, observational study. Dogs were recruited from 11 different veterinary clinical sites across the continental US from January 2021 to August 2021 via word-of-mouth, email communication to existing clientele, or social media advertising. Dogs were included if they met the following age and size criteria: ≥ 2 and < 6 years of age (young dogs) **or** ≥ 7 years of age (old dogs); <11.3 kg / <25 lbs (small dogs) **or** large > 22.7 kg / >50 lbs (large dogs). Age cutoffs were determined based on previous research norms to facilitate comparison between studies [34–36]. Similarly, consistent with previous research studies [37,38], weight cutoffs were selected based on AKC dog breed classification for small and large sized dogs [39]. All dogs had to be cooperative for study procedures without the use of injectable sedation. Accurate age was verified with a documented birth year on a breeding or a medical record from < 6 months of age.

**Fig. 1.**
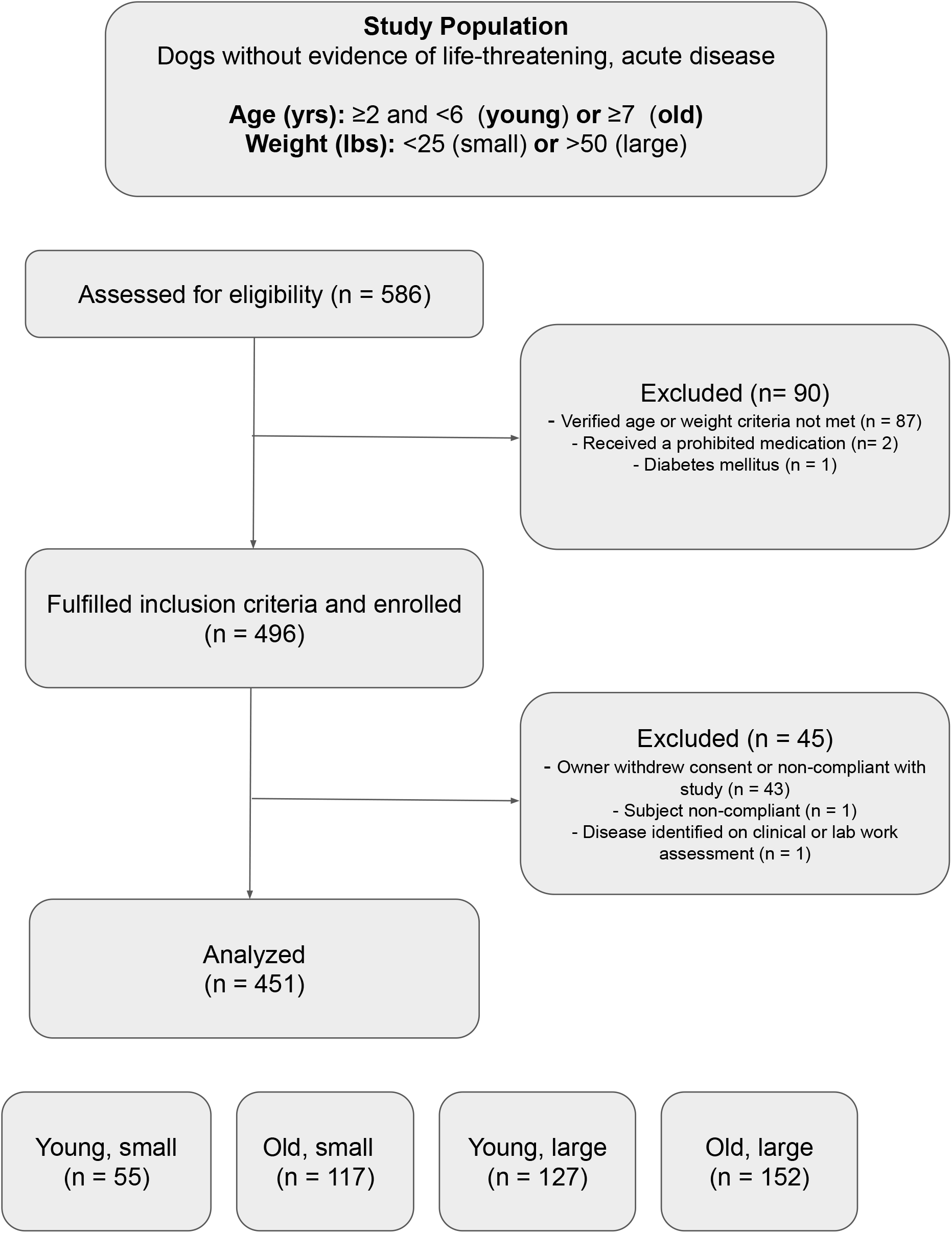
Five hundred and eighty-six dogs were screened for eligibility. Dogs without life-threatening acute disease, hypoadrenocorticism or diabetes were included if age was verified to be ≥2 and <6 years (young) or ≥7 years (old) and if body weight was < 25 lbs (small) or > 50 lbs (large). Of 586 dogs, 90 dogs were excluded primarily because age could not be verified or when age/weight criteria were not met (n = 87). Two dogs received a prohibited medication and 1 dog was excluded due to a history of diabetes mellitus. Four hundred and ninety-six met the inclusion criteria for enrollment of which 45 dogs were subsequently excluded due to owner non-compliance (n = 43), subject non-compliance (n = 1), or disease identified on evaluation of physical exam or bloodwork (n = 1). Four hundred and fifty-one dogs were analyzed of which 55 were young and small, 117 old and small, 127 young and large, and 152 old and large

Dogs were excluded from the study if they had a history or physical examination suggestive of severe illness such as recent dramatic weight loss (≥ 10% of baseline) within the previous two weeks, severe dehydration, pale or white mucous membranes, labored breathing, or evidence of fever or hypothermia (rectal temperature > 103.5 or < 99 °F). Dogs were excluded if they were already enrolled in an ongoing clinical trial evaluating a pharmaceutical medication, nutraceutical, or supplement or if the dog had undergone a surgical or anesthetic procedure within 14 days of Day 0. In order to prevent confounding with future analysis of blood based biomarkers (not reported here), dogs were also excluded if they had previously been diagnosed with diabetes mellitus or hypoadrenocorticism or if they received corticosteroids within 14 days of the Day 0 study visit or if they had ever received metformin.

### Study Procedures

Owner Informed Consent (OIC) was obtained electronically before or on the day of the study visit (Day 0). The VetMetrica HRQL survey was completed on Day 0 before the dog was examined by a study veterinarian or up to 3 days prior. On Day 0, the Investigator or study veterinarian obtained medical history, performed a physical exam, and completed the CFI and evaluated final eligibility based on inclusion/exclusion criteria. Investigators were instructed to collect blood and urine samples from dogs fasted overnight or for at least 6 hours for comprehensive CBC, serum biochemical profile, urinalysis, and total thyroid hormone (T4). Duration of fasting was not verified. Investigators and study veterinarians reviewed all clinical laboratory results to finalize the CFI and sign off on final inclusion/exclusion criteria after Day 0. The schedule of events is detailed in **Table 1**.

**Table 1.** Schedule of events.

| **<u>Event</u>** | **<u>Day –</u>**<br>**<u>X<sup>a</sup></u>** | **<u>Day -3 to</u>**<br>**<u>Day 0<sup>b</sup></u>** | **<u>Day 0</u>** | **<u>Post</u>**<br>**<u>Day 0</u>** |
| --- | --- | --- | --- | --- |
| Screen for Inclusion/Exclusion Criteria | ✓ |  |  |  |
| Obtain Owner Informed Consent (OIC)<sup>a</sup> | ✓ |  |  |  |
| Health-Related Quality of Life (HRQL) *Owner Assessment<sup>b</sup>* |  | ✓ |  |  |
| Review Medical History and Signalment |  |  | ✓ |  |
| Record Concomitant Medications |  |  | ✓ |  |
| Physical Examination (PE) |  |  | ✓ |  |
| Body Condition Score and Muscle Condition Score |  |  | ✓ |  |
| Canine Frailty Index<br>(Investigator or Study Veterinarian Assessment)<sup>c</sup> |  |  | ✓ | ✓ |
| Review Inclusion/Exclusion Criteria |  |  | ✓ | ✓ |
| Hematology and Biochemistry |  |  | ✓ |  |
| T4 |  |  | ✓ |  |
| Urinalysis |  |  | ✓ |  |
| Study Exit<sup>d</sup> |  |  |  | ✓ |
| <sup>a</sup> The OIC was completed any time prior to or on day 0.<br><sup>b</sup> The HRQL was completed on day 0 or within 3 days prior to day 0.<br><sup>c</sup> The Canine Frailty Index was completed based on the dog's history, PE, and clinical pathology collected at day 0. Therefore, the form may have been completed on day 0 or after.<br><sup>d</sup> The Study Exit Form was completed in the Electronic Data Capture (EDC) once the laboratory results were reviewed. |  |  |  |  |

### Vetmetrica HRQL

Dog owners were registered in the NewMetrica system and instructed to complete the Vetmetrica HRQL survey prior to Day 0. If the survey was not completed before Day 0, owners were required to complete the HRQL survey on Day 0 at the clinical site before the veterinary examination. The electronic survey comprised of 22 psychometrically validated questions to capture four domains of quality of life – Energetic/Enthusiastic (E/E), Happy/Content (H/C), Active/Comfortable (A/C), and Calm/Relaxed (C/R). Each question consisted of a descriptor and was rated on a 7-point Likert scale, with a minimum of 0 (“not at all”) and a maximum of 6 (“could not be more”). The results were calculated and scored using proprietary algorithms to provide raw scores for each of the four quality of life domains. Higher numbers indicate better owner-perceived health-related quality of life.

### Canine Frailty Index

The Investigator or study veterinarian at each clinical site completed the Canine Frailty Index. Higher CFI scores indicated a greater number of health deficits (higher frailty) as recorded on the 33 item clinical questionnaire. Each item was scored as 0 for “absent”, 0.5 for “mild”, and 1 for “present” or “severe.” CFI scores were calculated by dividing the total sum of item scores by 33, with a minimum possible score of 0 and maximum of 1. The Investigator or study veterinarian completed the initial part of the assessment on day 0 based on the dog’s medical history and PE. The remainder of the assessment was completed after the Investigator or study veterinarian reviewed the clinical pathology results from the samples obtained on day 0.

### Statistical Methods and Analysis

Each individual dog served as the experimental unit. Missing data was not imputed and only observed data was included in statistical analyses.

Descriptive statistics (mean, standard deviation, median, interquartile range (IQR), and minimum and maximum) of demographic features (age, sex, and percentages of pure or mixed breed dogs) and key outcome variables (CFI score, total HRQL, and individual HRQL domain scores) were summarized for the entire dataset as well as by age-size group stratifications (young, small; old, small; young, large; old, large). CFI scores range from 0 to 1, with 1 indicating most frail. Individual HRQL domain raw scores range from 0-6, with larger scores indicating higher quality of life on that domain.

To ease interpretation, raw scores from the four domains (Energetic/Enthusiastic (E/E), Happy/Content (H/C), Active/Comfortable (A/C), and Calm/Relaxed (C/R)) were aggregated into a single score reflecting the total sum of scores across these domains (referred to as “total HRQL”). Following Davies [40], the four domains were first summed together, normalized by applying a logit transformation, standardized, and finally rescaled so that the distribution of the total scores range from 0 - 100 with mean 50 and standard deviation of 10. This total HRQL score reflects the overall quality of life for a given dog, where higher scores indicate better quality of life.

Differences in CFI, total HRQL and domain scores across age-size groups were assessed using one-way ANOVA with pairwise two-sample t-tests, adjusting for multiple comparisons using the Holm [41] method. If assumptions of ANOVA were violated, then nonparametric Kruskal-Wallis tests followed by pairwise comparisons with Wilcoxon rank sum tests were used.

Multiple linear regression modeling was used to explore the effects of age and body size on CFI, total HRQL, and HRQL domain scores. Due to study recruitment procedures, systematic gaps across continuous variables age and weight were present, precluding the ability to fit standard linear models testing interactions of weight and age. Therefore, to assess age-related rates of change in CFI and HRQL domain scores across age-size groups, we fit the following model:

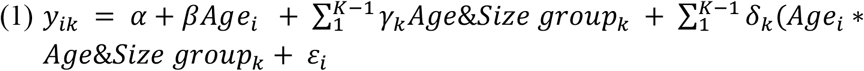

where *i* = 1,…, N dogs, *k* = 1, …, 4 age-size groups indicator variables with small, young dogs as the referent group. Estimated marginal means of linear trends (i.e., slopes for each age-size group) were estimated from this model and post-hoc linear contrast statements were then used to compare rates of change in CFI, total HRQL, and domain scores across the different age-size groups.

Simple and multivariate linear regression was used to assess the relationship between CFI score and total HRQL score and individual HRQL domain scores. Simple linear regression treated CFI score as the explanatory variable and HRQL total and domain scores as the outcomes. To explore if CFI score was associated with total HRQL and HRQL domain scores above and beyond effects of age and size, age and size-age group variables were also added into the model.

An exploratory mediation analysis was also performed with the most prevalent CFI items to identify the potential role of individual CFI deficits as mediators in any interaction effects of age and size group on CFI and HRQL domain scores. CFI items were first dichotomized as absence (“No”) = 0 and presence (“Yes”, “Mild”, or “Severe”) = 1. Relative prevalence for each item was reported across age-size groups. Multivariate logistic regression was then used to identify if age and size groups significantly interacted to predict individual CFI deficits. Deficits for which age and size group significantly interacted were deemed candidate mediators. Finally, candidate mediators were added into the interaction model (equation 1) to evaluate if the presence of each deficit mediated any interactions between age and size **(Fig. 2)**.

**Fig. 2.**
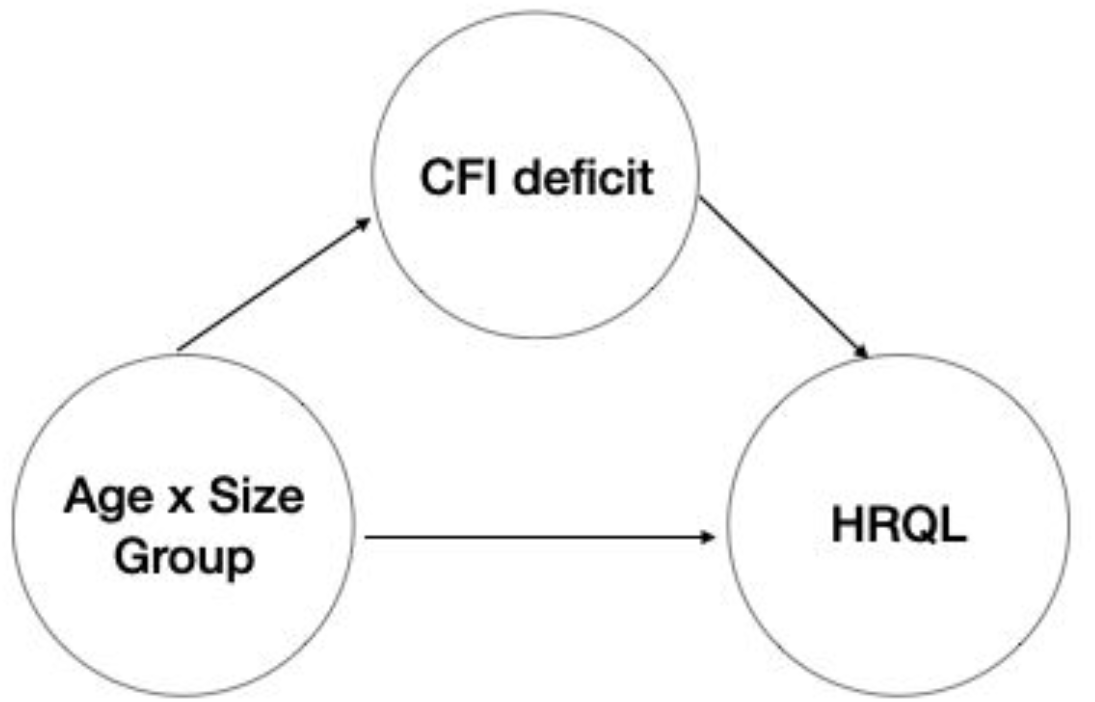
DAG (directed acyclic graph) represents the mediation analysis performed to evaluate whether individual CFI (Canine Frailty Index) deficits mediate the effect of age-size group on HRQL (Health Related Quality of Life) score

All regression models were tested for normality assumptions. All hypothesis tests of significance were performed at alpha = 0.05, two-sided, unless otherwise stated. All analyses and data presentations were performed using R version 4.12 [42].

## Results

### Subject Demographics

Following screening, 451 eligible adult dogs were enrolled of which 46.3% were female and 53.7% male. The mean age of dogs was 7.72 years (range = 2.0 to 17.7, SD = 3.8). Mean weight was 24.0 kg (range 1.3-80, SD 15). The majority (91.1%) of dogs were neutered. The evaluable population consisted of 43.6% mixed breeds and 56.4% purebred dogs. The demographics of the evaluated subjects in each of the age and size groups are presented in **Table 2**. A table of enrolled breeds <11.3 kg / <25 lbs (small dogs) or > 22.7 kg / >50 lbs (large dogs) is presented in Supp Table 1, Table 2, respectively.

**Table 2.**
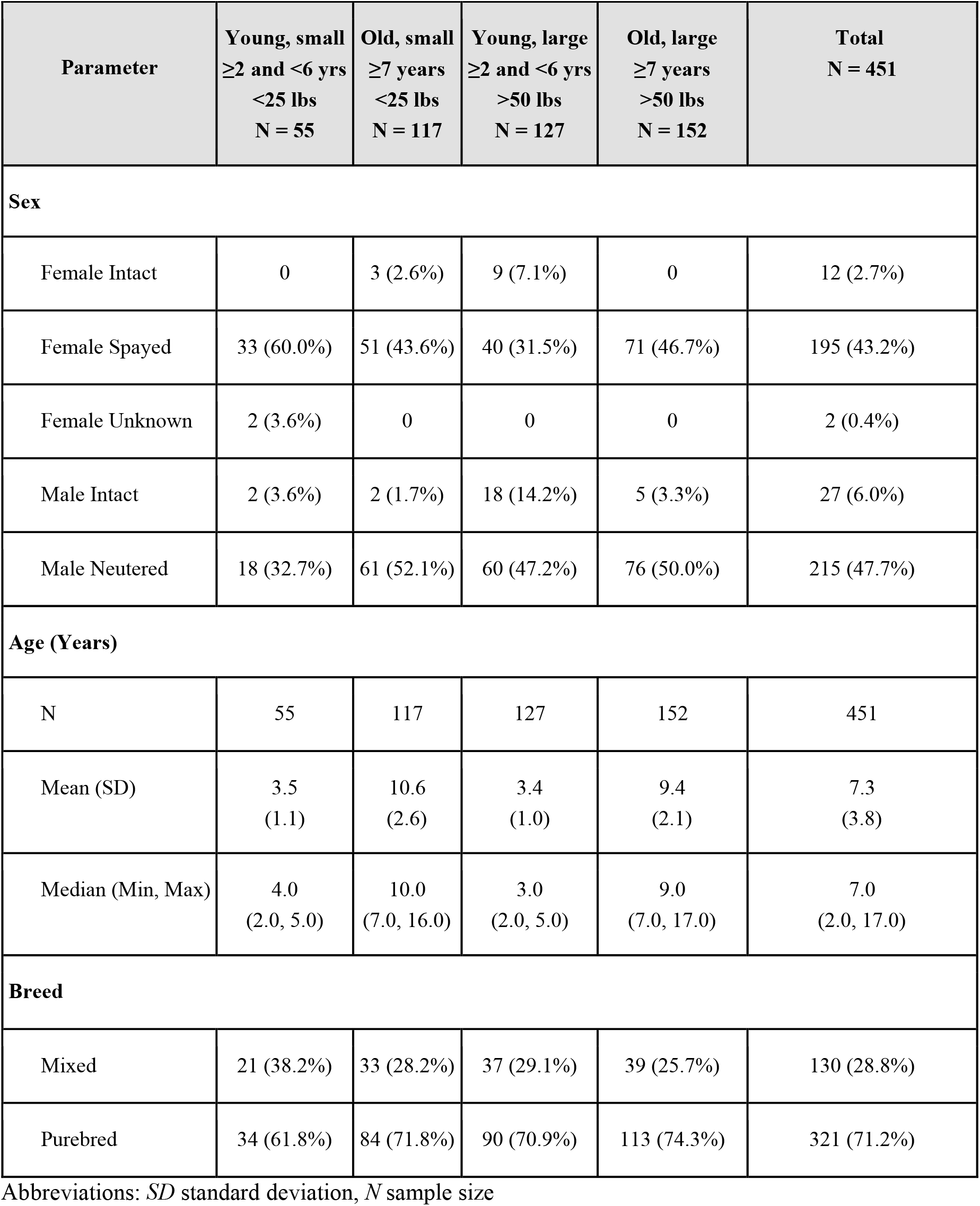
Demographics of study population with age, sex, and breed information in 4 age-size groups (young and small, old and small, young and large, old and large)

Of the total population (n = 451), 450 dogs had CFI scores and 448 dogs had VetMetrica HRQL domain scores (E/E, A/C, H/C, and C/R). One dog with the maximum observed CFI score of 0.7 was excluded upon confirmation that the CFI form had been completed erroneously by one of the study investigators. A total of 3 subjects were excluded from HRQL score analyses for failing to complete the HRQL on or prior to Day 0 (n = 1) or for refusing permission to use these data (n = 2).

### Canine Frailty Index (CFI) Scores increase with age

#### Wider distribution of scores in old versus young dogs

The distribution of CFI scores was wider in old dogs compared to young dogs. The distribution of the CFI scores in each age-size group was non-normal with positive skew. CFI scores in the evaluable population of dogs (n = 450) ranged from 0.0 to 0.59, with a median of 0.03, and IQR of 0.09. IQRs of CFI scores in old dog groups were larger than that of young dog groups (**Table 3**).

**Table 3.**
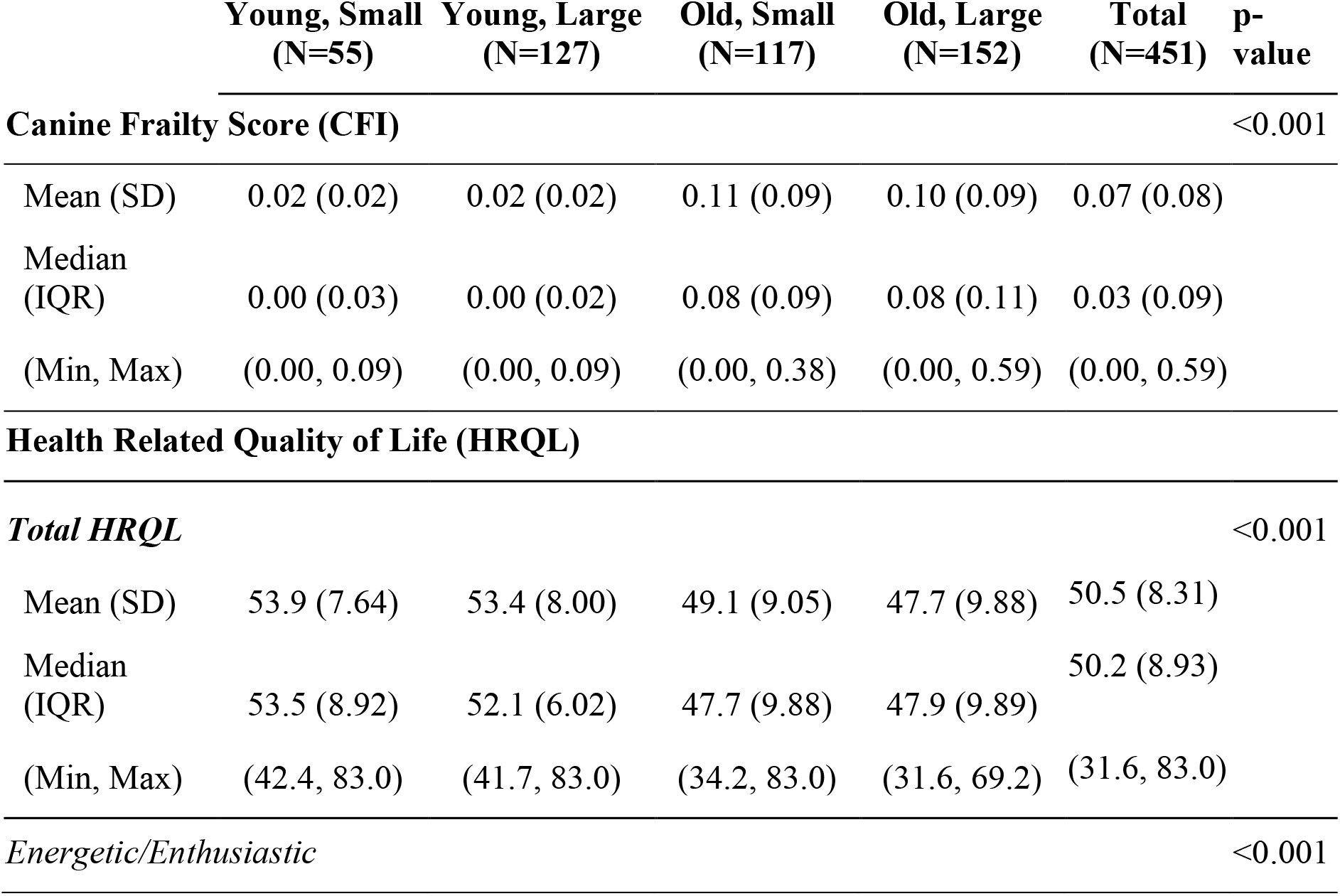
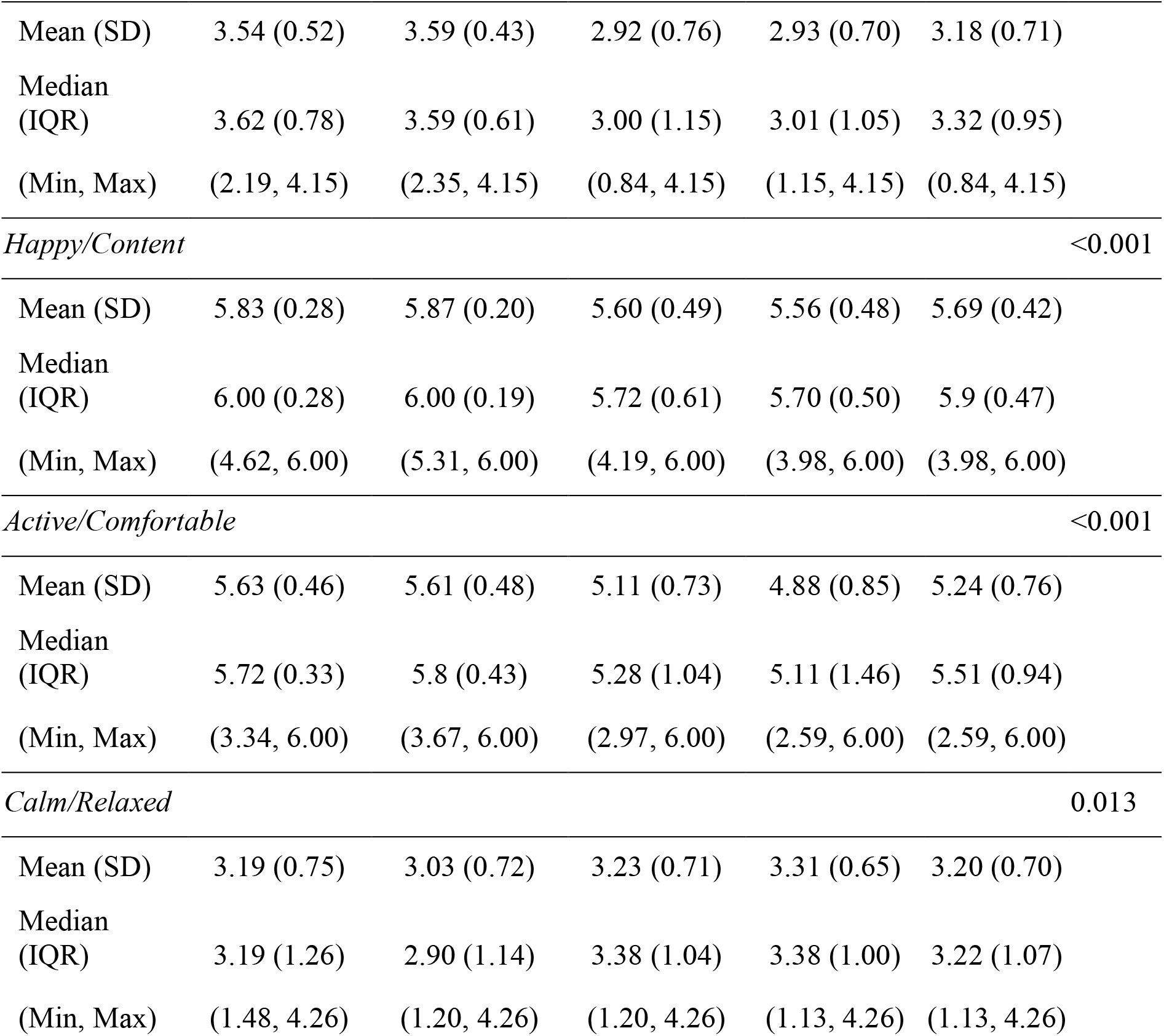
Canine Frailty Index (CFI) and Health Related Quality of Life (HRQL) Scores across age-size groups with mean, standard deviation, interquartile range, range (minimum, maximum), and p-values for Kruskal-Wallis Tests showing significant differences across age-size groups.

#### Higher CFI scores in old versus young dogs

Old dogs had significantly higher CFI scores compared to young dogs. A Kruskal-Wallis test showed a significant difference in CFI scores across age-size groups (H(3)= 182.87, p < 0.001) **(Table 3)**. Pairwise Wilcoxon-rank sum tests demonstrated that CFI scores were significantly higher in old dogs compared to young dogs, in both large (p-adjusted < 0.001) and small (p-adjusted < 0.001) dogs (**Fig. 3**).

**Fig. 3.**
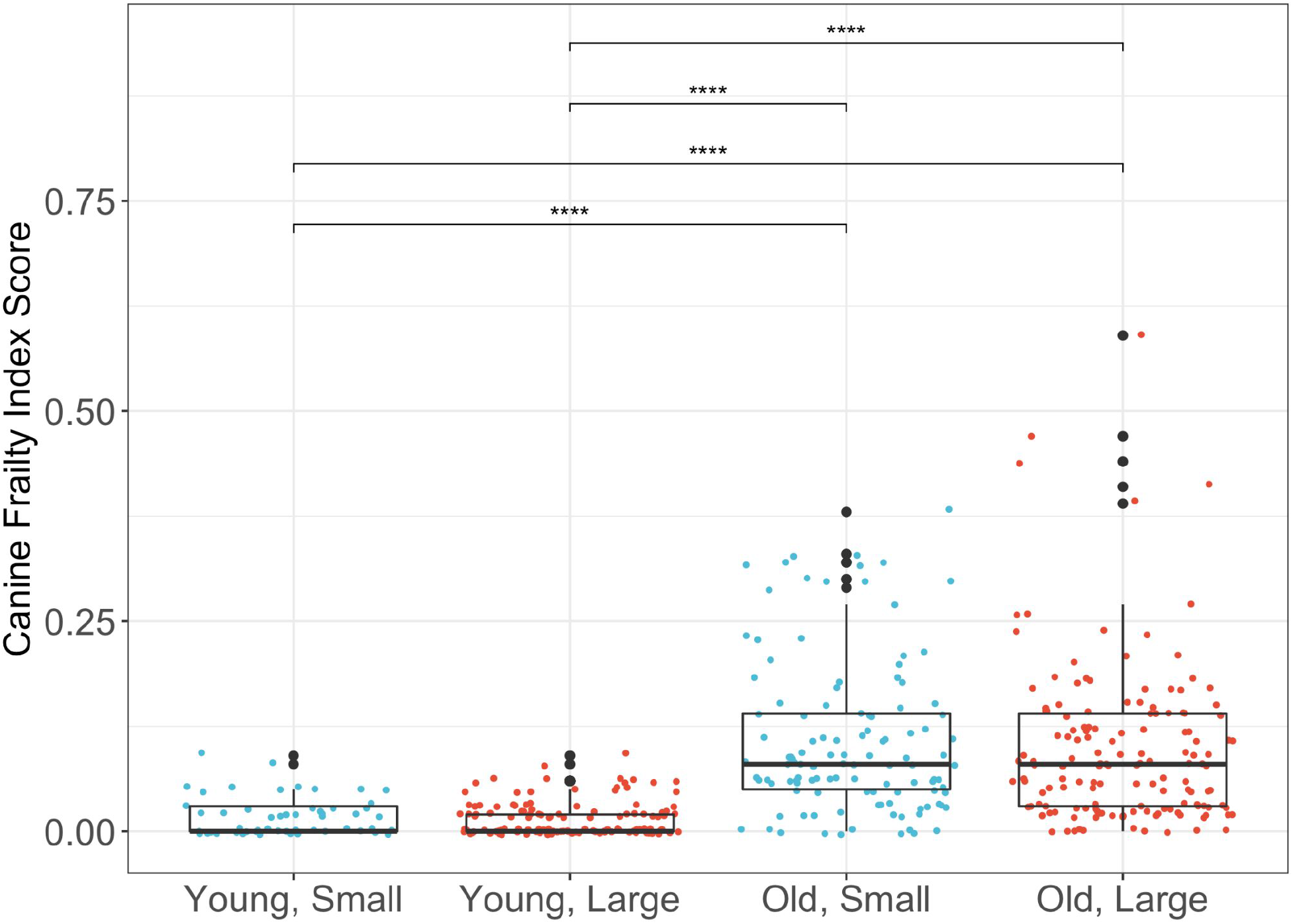
Canine frailty index scores in the different age-size groups (young and small, old and small, young and large, old and large) are shown with box and whisker plots. Young dogs were ≥ 2 and <6 years, old dogs were ≥7 years, small dogs were < 25 lbs and large dogs were > 50 lbs. The median is the line in the center of the box and the X represents the mean. The vertical line at the bottom of the box denotes the first quartile and the vertical line extending from the top of the box denotes the third quartile. Pairwise differences between young and old groups of corresponding size were compared with Wilcoxon rank sum tests. **** denotes p value <0.0001

#### CFI scores increased with age in older dogs but not younger dogs

Scatter plots showed trends of increasing CFI score with age (**Fig. 4**). Multiple linear regression analysis confirmed that CFI scores significantly increased with age in old dogs (old and small: *β*=0.02, p < 0.001; old and large: *β*=0.038, p < 0.001), but not in young dogs (young and small: *β*=0.00, p = 0.503; young and large: *β*=0.00, p = 0.383) **(Supplemental Table 3)**. Within the old and young age groups, old dogs had faster age-related increases in CFI score compared to young dogs (*β*_old_-*β*_young_=0.04, p < 0.001). While an overall interaction effect between age and age-size group was observed (F(3)= 9.49, p < 0.001), there were no size-based differences in age-related increases in CFI scores within old (*β*_old,large_-*β*_old, small_= 0.00, p = 0.278) or young dogs (*β*_young, large_-*β*_young, small_= 0.00, p = 0.944) **(Table 4)**.

**Fig. 4.**
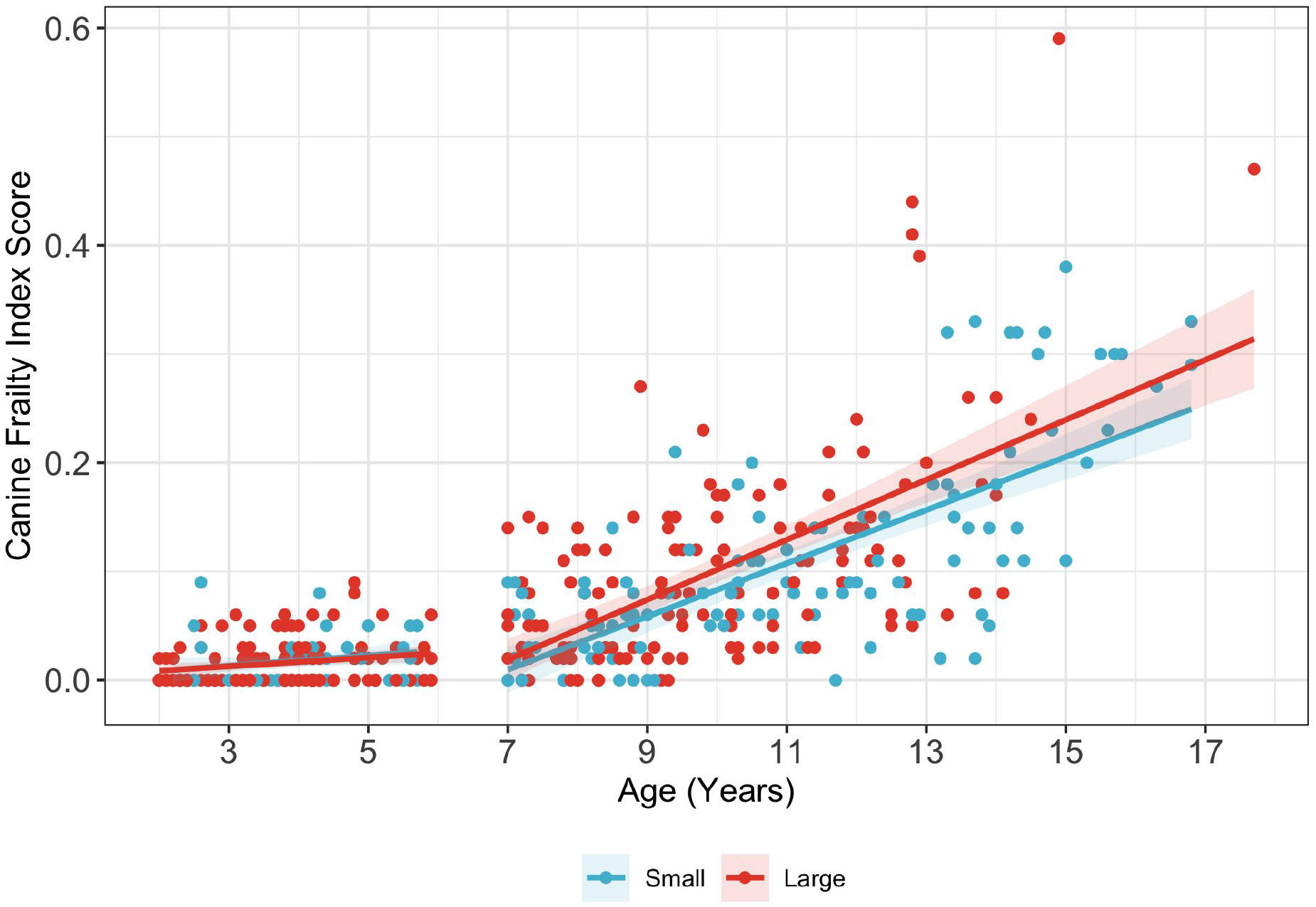
Canine frailty index (CFI) scores in the different age-size groups are depicted in scatter plots with a fitted line. Red dots denote the small dogs < 25 lbs and blue dots denote the large dogs > 50 lbs. Red and blue shading show the confidence intervals

**Table 4.** Linear contrast estimates comparing differences in age trends of Canine Frailty Index (CFI) and Health Related Quality of Life (HRQL) between age and age-size groups.

|  | Comparison | Difference in Age Effect (95% CI) | Comparison p-value<sup>a</sup> | Overall interaction effect p-value<sup>b</sup> |
| --- | --- | --- | --- | --- |
| **Canine Frailty Index (CFI)** |  |  |  |  |
|  | Old vs Young | 0.04 (0.03, 0.06) | **<0.001** | **<0.001** |
|  | Old Dogs: Large vs Small | 0.00 (-0.00, 0.01) | 0.278 |  |
|  | Young Dogs: Large vs Small | -0.00 (-0.02, 0.02) | 0.944 |  |
| **Health Related Quality of Life (HRQL)** |  |  |  |  |
| **Total HRQL** | Old vs Young | -1.37 (-3.71, 0.03) | 0.250 | 0.183 |
|  | Old Dogs: Large vs Small | 0.23 (-0.53, 0.99) | 0.554 |  |
|  | Young Dogs: Large vs Small | 1.38 (-0.84, 3.59) | 0.223 |  |
| Energetic/Enthusiastic | Old vs Young | -0.15 (-0.34, 0.03) | 0.106 | 0.124 |
|  | Old Dogs: Large vs Small | 0.02 (-0.04, 0.08) | 0.613 |  |
|  | Young Dogs: Large vs Small | 0.09 (-0.09, 0.26) | 0.332 |  |
| Happy/Content | Old vs Young | -0.14 (-0.26, -0.02) | 0.020 | 0.130 |
|  | Old Dogs: Large vs Small | -0.01 (-0.04, 0.03) | 0.774 |  |
|  | Young Dogs:<br>Large vs Small | -0.03 (-0.14, 0.09) | 0.624 |  |
| Active/Comfortable | Old vs Young | -0.27 (-0.47, -<br>0.08) | **0.006** | **<0.001** |
|  | Old Dogs: Large<br>vs Small | -0.07 (-0.14, -<br>0.01) | **0.022** |  |
|  | Young Dogs:<br>Large vs Small | 0.12 (-0.06, 0.30) | 0.201 |  |
| Calm/Relaxed | Old vs Young | -0.00 (-0.22, 0.21) | 0.971 | 0.987 |
|  | Old Dogs: Large<br>vs Small | 0.01 (-0.06, 0.08) | 0.739 |  |
|  | Young Dogs:<br>Large vs Small | -0.02 (-0.22, 0.19) | 0.872 |  |

Abbreviations: *CFI* Canine Frailty Index, *HRQL* Health Related Quality of Life, *CI* Confidence Interval <sup>a</sup>Comparison p-value indicates that contrasts are significantly different from zero when $p < 0.05$ (bolded). <sup>b</sup>Overall interaction effect corresponds to the likelihood ratio test for the interaction between age and age-size group. $p < 0.05$ are significant and bolded.

### HRQL Total & Domain Scores decrease with age

#### Wider distribution in scores in old versus young dogs

Older dogs had a wider distribution (higher IQR) of total and HRQL domain-specific scores than younger dogs, indicating more observed variability within the older population. Ranges, means, SDs, medians and IQRs for total and all individual HRQL domain scores are presented in **Table 3**. For individual HRQL scores, the same pattern across age bins was observed in all domains except C/R. Older dogs showed lower median HRQL scores than younger dogs across all domains, except C/R, in which older dogs scored higher **(Table 3)**.

#### Higher HRQL scores in old versus young dogs

Older dogs had significantly lower total HRQL scores compared to younger dogs. A Kruskal-Wallis test showed a significant difference in total HRQL scores between age-size groups (H(3)= 48.97, p < 0.001)) **(Table 3)**. Pairwise Wilcoxon rank sum tests demonstrated that older dogs had significantly lower total HRQL scores, compared to younger dogs, in both large and small dogs (**Fig. 5a**). The same pattern was observed for each individual HRQL domain score, where a Kruskal-Wallis test showed a significant difference between age-size groups (E/E: H(3)= 91.44, p < 0.001, H/C: H(3)= 54.00, p < 0.001, C/R: H(3)= 10.75, p = 0.013, A/C: H(3)= 94.56, p < 0.001). Pairwise Wilcoxon rank sum tests demonstrated that younger dogs had significantly higher E/E, H/C, and A/C scores compared to older dogs, regardless of size. In contrast, a size effect was only seen within the C/R (calm/relaxed) domain where only old, large dogs showed significantly higher C/R scores than young, large dogs (p-adjusted = 0.007, **Fig. 5b**), whereas there was no difference between old, small and young, small dogs.

**Fig. 5.**
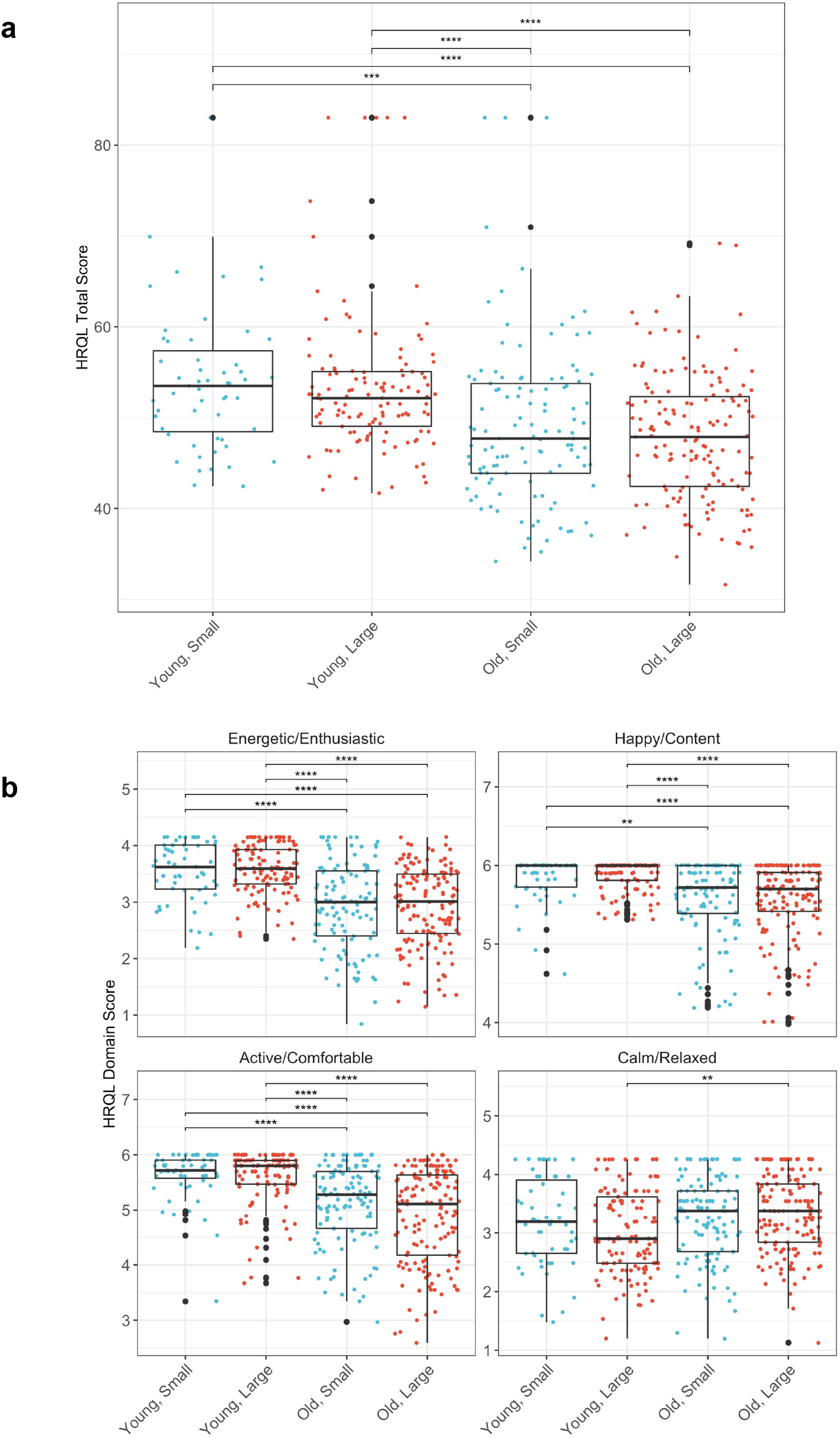
a. Total Health related quality of life (HRQL) scores are shown with box and whisker plots across age-size groups (young and small, old and small, young and large, old and large). Young dogs were ≥ 2 and <6 yrs, old dogs were ≥7 years, small dogs were < 25 lbs and large dogs were > 50 lbs. The median is the line in the center of the box and the X represents the mean. The vertical line at the bottom of the box denotes the first quartile and the vertical line extending from the top of the box denotes the third quartile. Pairwise differences between young and old groups of corresponding size were compared with Wilcoxon rank sum tests. ** p <0.01; **** p < 0.0001 b. Individual domain HRQL scores (Energetic/Enthusiastic, Happy/Content, Active/Comfortable, and Calm/Relaxed) are shown with box and whisker plots across age-size groups (young and small, old and small, young and large, old and large). Young dogs were ≥ 2 and <6 yrs, old dogs were ≥7 years, small dogs were < 25 lbs and large dogs were > 50 lbs. The median is the line in the center of the box and the X represents the mean. The vertical line at the bottom of the box denotes the first quartile and the vertical line extending from the top of the box denotes the third quartile. Pairwise differences between young and old groups of corresponding size were compared with Wilcoxon rank sum tests. ** p <0.01; **** p < 0.0001

#### HRQL scores declined with age in old dogs but not young dogs

Scatter plots showed trends of decreasing total and individual HRQL scores E/E, H/C, and A/C in old dogs, but not in young dogs. (**Fig. 6**). Multiple linear regression model results confirmed that old dogs showed significant age-related declines in total HRQL scores (old and small: *β*=-1.5, p <0.001, old and large: *β*=-1.3, p < 0.001) that were not observed in the young dogs. Individual HRQL domain scores, except for C/R, also showed age-related decline in old dogs but not in young dogs: E/E (old and small: *β*=-0.14, p <0.001, old and large: *β*=-0.12, p < 0.001), H/C (old and small: *β*=-0.06, p <0.001, old and large: *β*=-0.07, p < 0.001), and A/C (old and small: *β*=-0.13, p <0.001, old and large: *β*=-0.20, p < 0.001) (**Supplemental Table 3**).

**Fig. 6.**
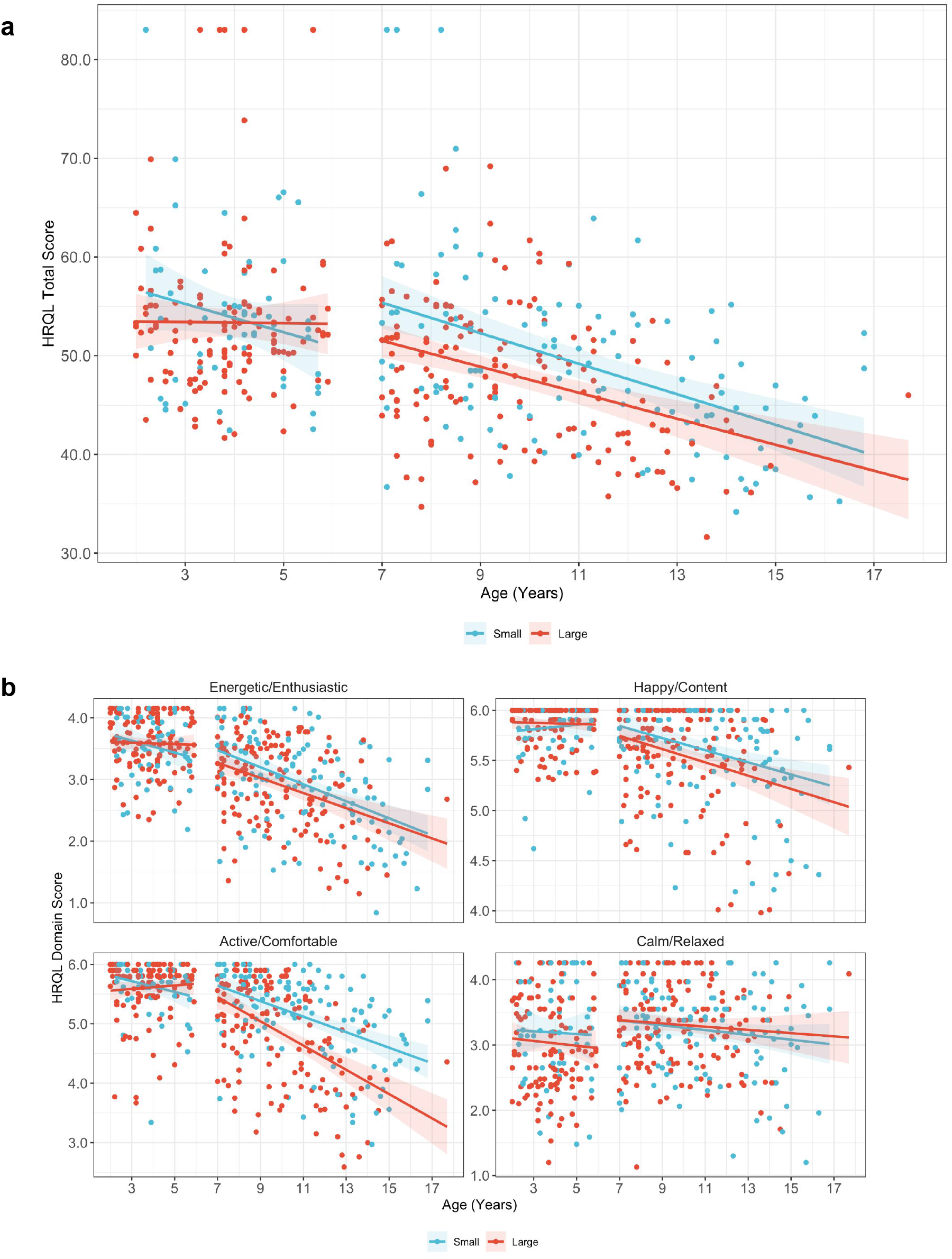
a. Total Health Related Quality of Life (HRQL) scores in the different age- and size groups are depicted in scatter plots with a fitted line for each domain. Red dots denote the small dogs < 25lbs and blue dots denote the large dogs > 50 lbs. Red and blue shading show 95% confidence intervals b. Individual domain Health Related Quality of Life (HRQL) scores in the different age- and size groups are depicted in scatter plots with a fitted line. Red dots denote the small dogs < 25lbs and blue dots denote the large dogs > 50 lbs. Red and blue shading show 95% confidence intervals

#### Faster age-related decline in A/C scores in older, larger dogs

Likelihood ratio tests of multiple linear regression models indicated a significant interaction effect between age and age-size group for individual A/C HRQL scores only (p<0.001), but not for other domains or total HRQL score **(Supplemental Table 3)**. Linear contrast statements comparing age-related changes in HRQL between age-size groups **(Table 4)** revealed that old dogs had significantly faster age-related declines in A/C scores compared to that of young dogs (*β*_old_-*β*_young_= -0.27, p = 0.006). Within old dogs, large dogs showed more rapid age-related decline in A/C scores compared to small dogs (*β*_old,large_-*β*_old, small_= -0.07, p = 0.022).

### Clinically-assessed CFI is inversely related to owner-assessed total HRQL and HRQL domain scores after adjusting for age

CFI score and total HRQL score were inversely related. Simple linear regression analyses demonstrated inverse linear relationships between CFI score and total HRQL score and all HRQL individual domains except for C/R scores (R^2^<0.01, *β* = -0.38, p = 0.344) **(Fig. 7)**. Multivariate regression models accounting for age and age-size group and their interaction showed that CFI score was significantly inversely associated with all HRQL scores (total and individual domains, even after adjusting for the effect of age: total HRQL score (*β* = -28.3, p < 0.001); E/E (*β* = -2.64, p < 0.001); H/C (*β* = -1.995, p < 0.001); A/C (*β* = -3.31, p < 0.001); C/R (*β* = -1.22, p = 0.042) **(Supplemental Table 5)**. Additionally, the previously observed finding of large dogs showing more rapid decline in A/C scores with age was still observed, even after adjusting for CFI score (*β*_old,large_-*β*_old, small_= -0.06, p = 0.042). Thus, we aimed to further explore the reason for this size difference in declining A/C score with age.

**Fig. 7.**
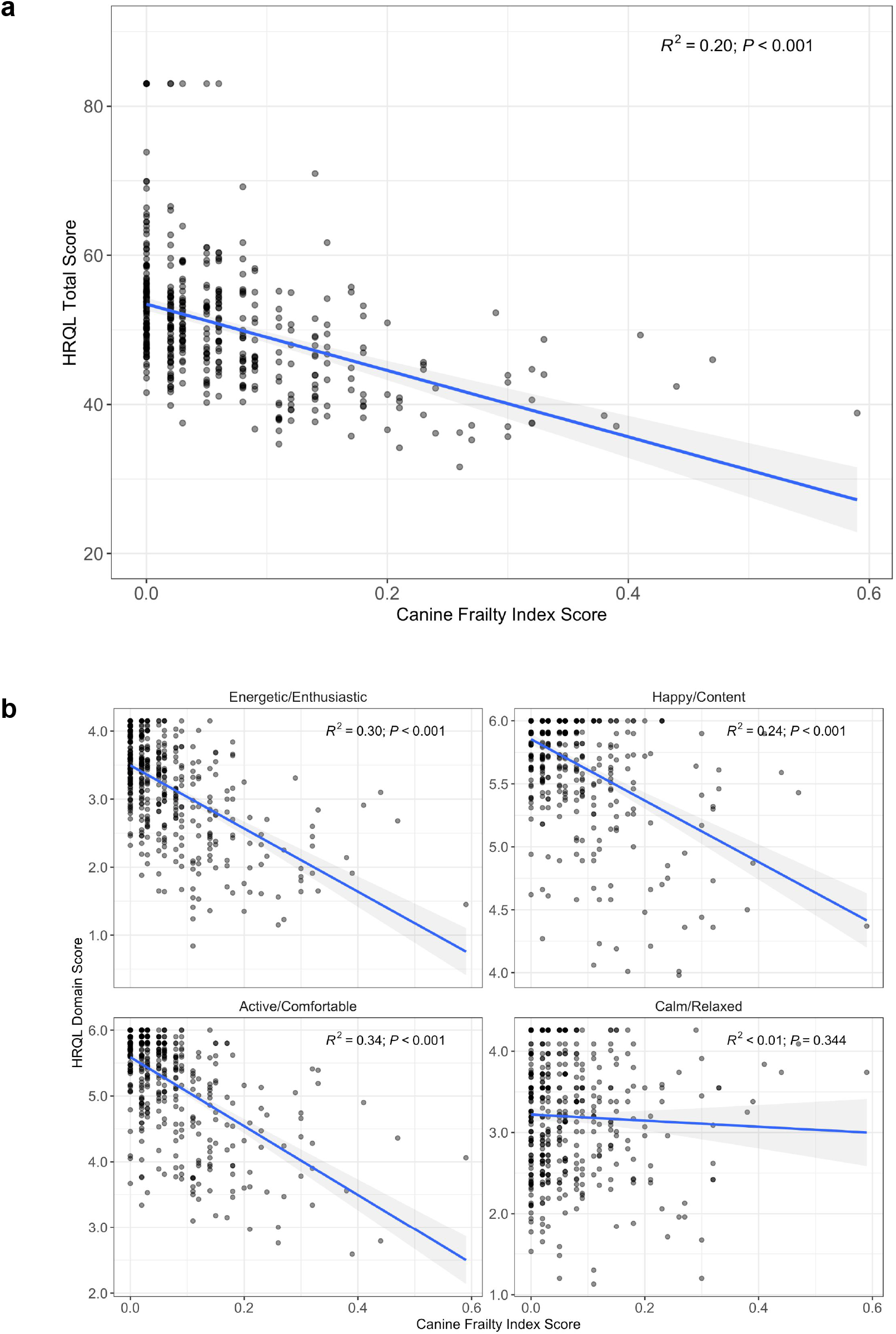
a. Scatter plots of total Health Related Quality of Life (HRQL) scores on the y axis against Canine Frailty Index (CFI) scores on the x axis with a fitted line. R^2^ is the correlation coefficient and the p-value indicates the significance of the correlation b. Scatter plots of individual domain HRQL scores on the y axis plotted against CFI scores on the x axis with a fitted line. R^2^ is the correlation coefficient and the p-value indicates the significance of the correlation

### Age-associated disease may mediate size differences in decline of A/C HRQL domain scores in old dogs

We evaluated clinical information collected from the CFI to see whether specific CFI clinical diseases or deficits might mediate the faster age-related decline in A/C scores in old, large dogs compared to old, small dogs. To do so, we calculated the relative prevalence across age-size groups, which yielded 12 CFI items with a total prevalence of at least 0.25. We found differences in relative prevalences in diseases between old, large and old, small dogs: oral disease was more common in small dogs, whereas osteoarthrosis (osteoarthritis), hair whitening, and decreased activity were more common in large dogs (**Fig. 8**). As outlined in the methods section, these were then individually tested as candidate mediators by examining the interaction effect of age and size group on these clinical deficits. Results of these logistic regression analyses indicated that size group (large or small) significantly interacted with age in explaining osteoarthritis (OA) prevalence, making it a candidate mediator (F(1) = 4.52, p=0.034) **(Supplemental Table 4)**. To complete our exploratory mediation analysis, OA was added into the full interaction model of A/C and was found to be significantly associated with reduced A/C scores (*β* = -0.50, p < 0.001). Its inclusion did not eliminate the significant interaction effect of age and age-size group on A/C (F(3)=3.52 p=0.015). OA, as opposed to other CFI items, may be a partial mediator for the faster A/C decline in old, large dogs compared to old, small dogs. **(Supplemental Table 4)**.

**Fig. 8.**
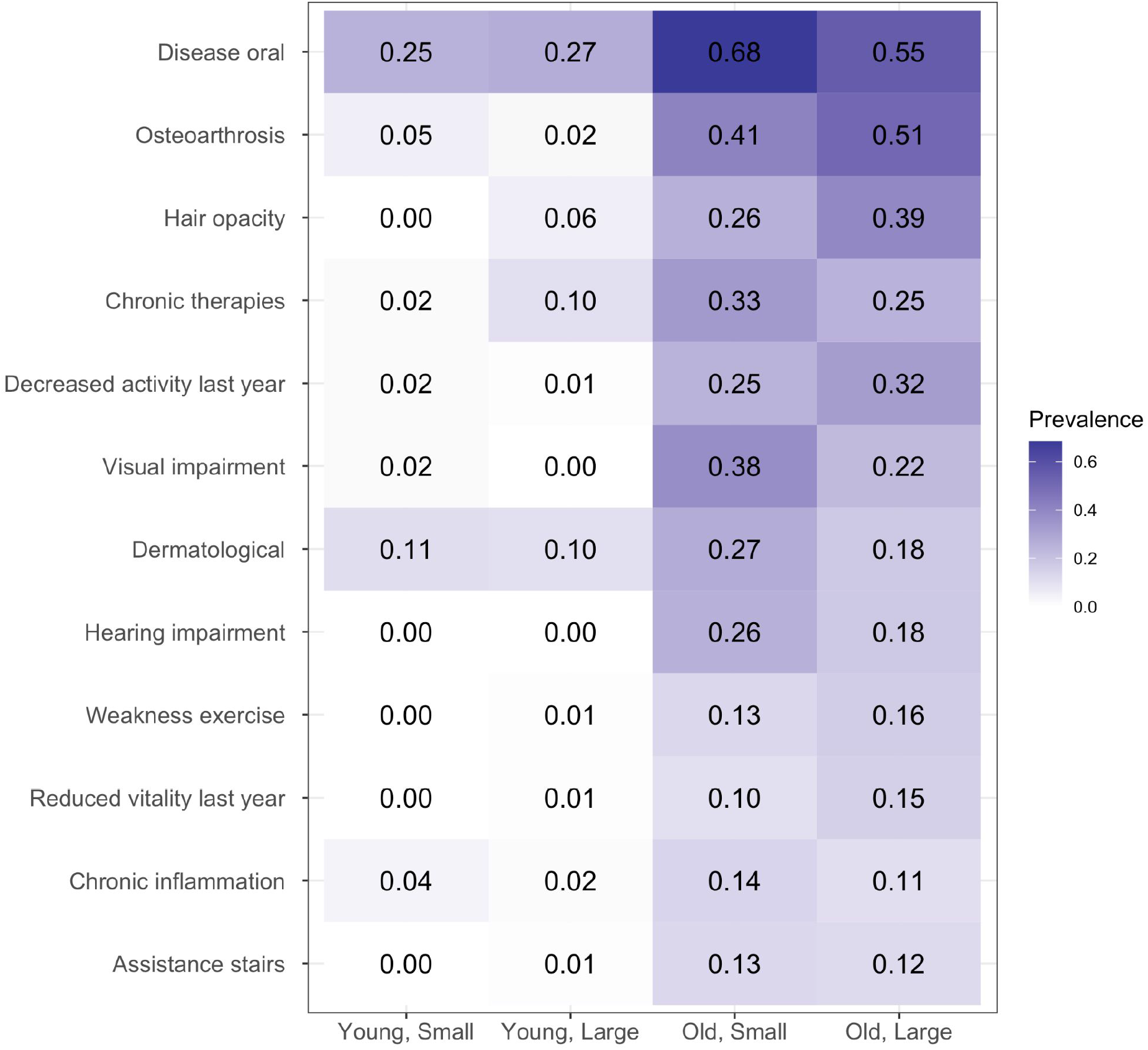
The most common Canine Frailty Index (CFI) Features in each of the age-size groups are listed with the heat map indicating darker colors of higher relative prevalence in this study population

## Discussion

As the companion dog emerges as a translational model for human aging and age-related disease, the development and validation of healthspan assessment tools in dogs is important for evaluating gerotherapeutics and advancing biogerontology research. This study aimed to assess two previously published instruments (Canine Frailty Index & NewMetrica’s HRQL) for measuring and relating two key components of healthspan in companion dogs: frailty and health-related quality of life. To assess the validity of the CFI and VetMetrica HRQL, we hypothesized that the CFI and VetMetrica HRQL instrument would show age group differences with increased frailty scores and decreased HRQL scores in old dogs compared to young dogs. We also hypothesized that owner-assessed HRQL scores would show an inverse relationship with clinician-assessed CFI scores. Finally, we used both the overall scores as well as more granular information collected from these two instruments to investigate relationships between frailty, HRQL, age as a continuous measure, and body size. We expected that frailty scores would increase with age while HRQL scores would decline with age. We also expected that size may have an effect on CFI or HRQL score, where larger body size would be associated with higher frailty score or lower HRQL scores.

As expected, results showed that CFI scores, a measure of clinically assessed frailty, were significantly higher in old (≥ 7 years) compared to young dogs (≤ 2 to < 6 years). Total HRQL domain scores as well as individual HRQL scores related to energy and enthusiasm (E/E), happiness and contentment (H/C), and physical activity and comfort (A/C) were significantly lower in old compared to young dogs. CFI and HRQL scores were inversely associated, even after adjusting for the effect of age. These results, demonstrating expected relationships between frailty and HRQL scores in old and young dogs, show that the CFI and VetMetrica HRQL can be used to assess age-related changes in frailty and health-related quality of life in companion dogs. Additionally, these tools allowed for the exploration of relationships between age, body size, owner-assessed HRQL, and clinically-assessed frailty and multimorbidity. Frailty score increased with age, and HRQL scores decreased with age in old dogs but not young dogs. While body size did not affect the age-associated increase in CFI score or decline in total HRQL score, our results demonstrated a more rapid owner-observed decline in A/C score decline in large, old dogs compared to small, old dogs. Because the CFI provided standardized information on clinically assessed disease and deficits, we were able to explore potential differences in age-associated morbidities, such as OA, in mediating this effect. In this way, the CFI and the VetMetrica HRQL can provide complementary readouts of age-associated declines and healthspan in companion dogs.

### Relationships between CFI scores, age, and size

Consistent with previous findings [21], a significant positive relationship was found between CFI score and age. The present study also found that increases in CFI scores were significantly greater with every additional year of age in dogs ≥ 7 years aged compared to dogs between 2 to 6 years. This implies that frailty increases as dogs age and accumulates more rapidly in dogs ≥ 7 years age. The study population in Banzato et al., which evaluated CFI scores in association with age, had an overall higher disease burden and dogs were enrolled from a single tertiary referral academic hospital in Italy. We found a similar rate of increase in CFI score per year (∼0.02 /yr) in our study population of dogs presenting to predominantly primary care practices throughout the United States. This illustrates that the CFI may be a broadly applicable tool that effectively measures increases in frailty in dogs with varying disease across a diverse set of veterinary investigators and clinical practices. As reported by Banzato and colleagues, CFI score may also be moderately predictive of short-term mortality based on previous findings [21], but survival was not assessed in this study. Assessing the relationship of CFI score with survival of dogs in this study through longitudinal follow up will be required to evaluate whether the CFI is a frailty tool predictive of mortality in dogs.

When accounting for age, CFI scores were not significantly higher in large dogs compared to small dogs in this study, which suggests that overall multimorbidity may not be higher in large dogs despite consistent evidence of their shortened lifespans [28,29,43,44]. Additionally, the slopes of increasing CFI score with age did not significantly differ between large and small dogs, suggesting that large dogs do not accumulate age-associated disease and functional deficits more rapidly. These results are consistent with findings from previous cross-sectional studies on size related differences in canine multimorbidity, which showed that neither body weight nor breed had an observed effect on the number of comorbid diseases [32] or on cognitive aging, where the trajectory of cognitive decline did not differ due to size or breed-average lifespan [33]. Of course, it is possible that the cross-sectional differences between large and small dogs found here do not reflect longitudinal changes in individual dogs over time. Ongoing large scale longitudinal cohort studies [15,45] characterizing the natural course of canine aging will be invaluable for providing additional datasets to address whether size mediates any relationships between healthspan and lifespan in companion dogs.

### Relationships between HRQL scores with CFI, age, and size

Measuring quality of life to integrate the human patient (or pet-owner) perspective in response to an intervention is an important efficacy readout implemented in human clinical practice and clinical trials [46–48]. Quality of life is a central component of the concept of healthspan and measuring the number of “healthy” or “good quality” years lived could be operationalized by using these psychometrically validated HRQL instruments. While the VetMetrica HRQL instrument has been psychometrically validated [24,25] and differentiates between healthy and ill dogs [40], this study is the first to assess age and size differences in HRQL in conjunction with a standardized assessment of frailty, which we believe to be critical components of healthspan.

In the current study, total HRQL score and all individual HRQL domain scores (E/E, H/C, A/C, C/R) showed significant inverse correlations with clinician assessed CFI scores after accounting for the effect of age. This provides evidence that frailty contributes to age-related decline in HRQL scores and that an owner assessment can capture the adverse impact of frailty on health-related quality of life. The association between CFI and HRQL score was weakest for the C/R domain perhaps because C/R domain scores are a better reflection of differences or changes in temperament rather than declining health status or frailty [25,40]. In this study, we aggregated raw individual domain scores to a normalized total HRQL score, which could simplify interpretation and improve usability for clinicians and owners. Total HRQL score did not show differences between size groups nor interaction effects between age and age-size group, lending support to use total HRQL score as a global measure of HRQL in dogs regardless of body size. However, multivariate regression analyses demonstrated significant interaction effects between age and age-size group for A/C domain scores, specifically in older dogs. This suggests that individual domains such as A/C may be more useful for research purposes by providing more granular information on size/breed-dependent differences in age-related declines of physical activity and comfort.

### Size-related differences in CFI deficit prevalence and the individual HRQL domain score A/C

The CFI not only demonstrated utility in standardizing and quantifying frailty, but also allowed us to explore qualitative differences in age-associated diseases between large and small dogs. Although this study did not observe any size-related differences in overall CFI scores, exploring size-related differences in specific CFI diseases and deficits were observed between small and large dogs. The most prevalent CFI items overall were more common in old compared to young dogs. Oral disease was very common in all four age-size groups, but was the most common in old, small dogs. Osteoarthrosis, hair whitening, decreased activity in the last year, weakness with exercise, reduced vitality in the past year were more common in large compared to small dogs. These differences were consistent with studies showing that small dogs have increased periodontal disease [49] and that large size is a risk factor for osteoarthritis [50,51]. Our exploratory mediation analysis allowed us to explore how size-related differences in age-associated disease may modulate the decline in A/C scores, introducing future hypotheses to formally test in future studies. As a clinically assessed frailty index, the CFI could be useful for global quantification of frailty applicable to any sized dog, while also uncovering breed and size differences in age-associated diseases.

### Limitations & future directions

There are several limitations to consider in this study. Firstly, due to its cross-sectional design, this study does not provide information on the longitudinal performance of the HRQL and CFI.

In the above analyses we did not control for sex, neuter status, or breed effects. Purebred dogs are documented to have shorter lifespans compared to mixed breeds and likely age differently from mixed breed dogs due to inbreeding depression [28,29,52]. Environmental and owner-influenced factors, such as exercise and diet were not considered in this study, which are potential confounding factors.

This study did not assess dogs between 6 and 7 years and dogs between 25-50 lbs/11.3 - 22.7 kgs. These age and weight cutoffs were chosen to differentiate between and compare groups, but precludes full characterization of the relationship between size and age on HRQL and CFI.

Additional selection bias may have occurred due to challenges in recruiting old and frail dogs. For instance, owners of dogs with higher multimorbidity and frailty are less likely to enroll in an observational study. This could have biased the study population towards old dogs with better health status than the population at large. In general, geriatric dogs (>12 years old) were not highly represented in this study population and small sized dogs are more highly represented at older ages.

Finally, both HRQL and CFI are subject to measurement bias. Despite undergoing psychometric validation, the VetMetrica HRQL is ultimately a subjective measurement that may be sensitive to response style effects such as the owner’s tendency to select extremes on the rating scale. Clear ceiling effects are observed in the HRQL data, suggesting a limited range of sensitivity to detect differences in HRQL at young ages. Ideally, HRQL instruments should be complemented by objective measures of the functional impact of aging. Similarly, floor effects are observed in the CFI data, suggesting limited sensitivity and interpretability at young ages when health deficits and diseases are uncommon.

While the CFI is veterinarian assessed, there are items of the CFI that may be vulnerable to noise and interobserver variability if investigators are not trained on how to assess and score items. In the present study, interobserver reliability was not formally assessed. However, the CFI was robust enough to show higher frailty with increasing age, at a similar rate found in Banzato et al., 2019, even although CFI was scored by a dozen study veterinarians and investigators across 11 different sites. Evaluating interobserver reliability in future studies will further strengthen the validity and utility of this tool.

## Conclusion

This 451 dog cross-sectional pilot study demonstrated the ability of two different tools (the Canine Frailty Index and the VetMetrica HRQL instrument) to identify age-related differences across a US companion dog population. CFI scores increased with older age while total HRQL score, as well as individual domain scores for E/E, H/C, and A/C decreased, confirming the expectation that frailty increases and health-related quality of life declines as dogs age. These trends were largely consistent across size groups, suggesting that dogs experience similar age-related changes in frailty regardless of body size. However, A/C scores decreased not only with age, but also with larger body size. Altogether, the results of this study lend support for the use of the Canine Frailty Index and VetMetrica HRQL in capturing two key components of healthspan: the increase in frailty and deterioration in quality of life with age. Characterizing the ability of the CFI and VetMetrica HRQL tools to effectively detect age and size-related differences in frailty and HRQL amongst companion dogs paves the way to utilize these tools in clinical and research settings to operationalize healthspan assessment in dogs and ultimately, evaluate novel gerotherapeutics that benefit both human and veterinary medicine.

## Acknowledgements

We thank all of our study participants and their dogs, and we acknowledge all of the work of our clinical site investigators and site personnel-without their efforts and contributions this study would not have been possible.

We would like to thank M. Kaeberlein, K. Creevy, D. Promislow, S. Niessen, K. Hamilton, N. Ehrhart, N. Olby, and M. Gruen for engaging in useful discussions regarding study design or results. We thank J. Reid and T. Banzato for discussion for implementation of the HRQL and CFI, respectively. We acknowledge M. Bell and A. Naka for guidance on statistical analysis; K. Greenwood & D. Juarez-Salinas for feedback on study design, final review, and proofreading; C. Lea Halioua-Haubold for final review; A. Hickson and T. L’Homme for contributions to initial study conduct and administration.

## Author Information

### Affiliations

Cellular Longevity Inc.

### Contributions

Conceptualization - FLC, ERR, MLF; Data Curation - FLC, JLG; Formal Analysis - JLG, FLC, JV, TVU; Investigation - FLC, BM, TAC, KMS, JA, SYW; Project admin - FLC, TAC, SYW; Writing- original draft- FLC, ERR, TVU; Writing- review and editing - FLC, TVU, MLF; Supervision – MLF

### Ethics Declaration

Authors declare IACUC approval (#VACLOY001CLDEFF1PILC) and owner informed consent was obtained for all study procedures.

