## Supplemental Tables for "Evaluating instruments for assessing healthspan: a multi-center cross-sectional study on health-related quality of life (HRQL) and frailty in the companion dog"

### Supplementary

**Supplementary Table 1**
**Enrolled breeds (<25 lbs)**

| Breed | Number of Dogs |
|---|---|
| Small mixed breed | 77 |
| Yorkshire Terrier | 18 |
| Shih Tzu | 17 |
| Dachshund | 16 |
| Chihuahua | 15 |
| Miniature Poodle | 14 |
| Pomeranian | 6 |
| Bichon Frise | 5 |
| Cavalier King Charles | 5 |
| Miniature Schnauzer | 5 |
| Papillon | 5 |
| Shetland Sheepdog | 5 |
| Pug | 4 |
| Beagle | 3 |
| Border Terrier | 3 |
| Cockapoo | 3 |
| French bulldog | 3 |
| Havanese | 3 |
| West Highland White Terrier | 3 |

**Supplementary Table 2**
**Enrolled breeds (>50 lbs)**

| Breed | Number of Dogs |
|---|---|
| Large mixed breed | 109 |
| Labrador Retriever | 60 |
| Golden Retriever | 33 |
| German Shepherd Dog | 17 |
| Boxer | 11 |
| Greyhound | 7 |
| Rhodesian Ridgeback | 7 |
| Bulldog | 6 |
| Doberman Pinscher | 6 |
| Rottweiler | 5 |
| Siberian Husky | 5 |
| Standard Poodle | 5 |
| Australian Shepherd | 4 |
| Irish Setter | 4 |
| Bernese Mountain Dog | 3 |
| Bloodhound | 3 |
| German Shorthaired Pointer | 3 |
| Goldendoodle | 3 |
| Great Dane | 3 |
| Mastiff | 3 |
| Viszla | 3 |
| Weimaraner | 3 |

#### Supplementary Table 3
#### Group age trend estimates for Health Related Quality of Life (HRQL) and Canine Frailty Index (CFI)

| | Age-Size Group | Estimated Effect of Age (95% CI) | p-value | Overall interaction effect p-value |
|---|---|---|---|---|
| **Canine Frailty Index (CFI)** | | | | |
| | Young, Small | 0.00 (-0.01, 0.02) | 0.503 | **<0.001** |
| | Young, Large | 0.00 (-0.01, 0.01) | 0.383 | |
| | Old, Small | 0.02 (0.02, 0.03) | **<0.001** | |
| | Old, Large | 0.03 (0.02, 0.03) | **<0.001** | |
| **Health Related Quality of Life (HRQL)** | | | | |
| ***Total HRQL*** | Young, Small | -1.43 (-3.28, 0.41) | 0.127 | 0.183 |
| | Young, Large | -0.06 (-1.28, 1.17) | 0.926 | |
| | Old, Small | -1.5 (-2.05, -1.04) | **<0.001** | |
| | Old, Large | -1.3 (-1.88, -0.75) | **<0.001** | |
| Energetic/Enthusiastic | Young, Small | -0.10 (-0.24, 0.05) | 0.187 | 0.124 |
| | Young, Large | -0.01 (-0.11, 0.08) | 0.816 | |
| | Old, Small | -0.14 (-0.18, -0.10) | **<0.001** | |
| | Old, Large | -0.12 (-0.17, -0.08) | **<0.001** | |
| Happy/Content | Young, Small | 0.02 (-0.07, 0.12) | 0.643 | 0.130 |
| | Young, Large | -0.01 (-0.07, 0.06) | 0.851 | |
| | Old, Small | -0.06 (-0.09, -0.03) | **<0.001** | |
| | Old, Large | -0.07 (-0.10, -0.04) | **<0.001** | |
| Active/Comfortable | Young, Small | -0.09 (-0.24, 0.06) | 0.246 | **<0.001** |
| | Young, Large | 0.03 (-0.07, 0.13) | 0.571 | |
| | Old, Small | -0.13 (-0.17, -0.09) | **<0.001** | |
| | Old, Large | -0.20 (-0.25, -0.16) | **<0.001** | |
| Calm/Relaxed | Young, Small | -0.02 (-0.19, 0.15) | 0.818 | 0.987 |
| | Young, Large | -0.04 (-0.15, 0.08) | 0.524 | |
| | Old, Small | -0.04 (-0.08, 0.01) | 0.128 | |
| | Old, Large | -0.02 (-0.08, 0.03) | 0.360 | |

#### Supplementary Table 4
#### Osteoarthritis mediation analysis

| Outcome | Covariate | Estimate (95% CI) | p-value | LRT p-value |
|---|---|---|---|---|
| Osteoarthritis (OA)[a.] | Age (years) | 1.53 (1.35, 1.77) | <0.001 | |
| | Size Group (Large) | 0.33 (0.04, 2.72) | 0.292 | |
| | Age*Size Group (Large) | 1.25 (1.02, 1.55) | 0.034 | 0.034 |
| HRQL A/C score | Age (years) | -0.07 (-0.22, 0.08) | 0.353 | |
| | Osteoarthritis (yes) | -0.50 (-0.66, -0.35) | <0.001 | |
| | Age-Size Group | | | |
| | Young, Small | ref | | |
| | Young, Large | -0.4 (-1.12, 0.31) | 0.266 | |
| | Old, Small | 0.30 (-0.46, 1.06) | 0.434 | |
| | Old, Large | 0.62 (-0.14, 1.37) | 0.108 | |
| | Age (years)*Age-Size Group | | | 0.015 |
| | Age*Young, Small | ref | | |
| | Age*Young, Large | 0.09 (-0.08, 0.27) | 0.293 | |
| | Age*Old, Small | -0.01 (-0.17, 0.14) | 0.862 | |
| | Age*Old, Large | -0.07 (-0.23, 0.08) | 0.339 | |

*OA* osteoarthritis, *ref* referent group, *LRT* Likelihood Ratio Test, *HRQL* Health-Related Quality of Life, *A/C* Active/Comfortable

a. Osteoarthritis modeled using logistic regression, all corresponding coefficient estimates are odds ratios

#### Supplementary Table 5
#### Multivariate regression of Canine Frality Index (CFI) on Health Related Quality of Life (HRQL) scores and corresponding contrasts

**Multivariate regression showing between CFI and HRQL total and Domain Scores**

| **Outcome** | **Estimated Effect of CFI (95% CI)** | **p-value** | **Overall interaction effect** |
|---|---|---|---|
| Total HRQL | -28.25 (-40.63, -15.86) | <0.001 | 0.443 |
| Energetic/Enthusiastic | -2.64 (-3.60, -1.68) | <0.001 | 0.441 |
| Happy/Content | -2.00 (-2.62, -1.37) | <0.001 | 0.814 |
| Active/Comfortable | -3.32 (-4.32, -2.31) | <0.001 | 0.031 |
| Calm/Relaxed | -1.22 (-2.4, -0.05) | 0.042 | 0.891 |

**Age trend estimates and contrasts for A/C after adjusting for CFI**

| **Age-Size Group** | **Estimated Effect of Age (95% CI)** | **p-value** |
|---|---|---|
| Young, Small | -0.07 (-0.22, 0.07) | 0.315 |
| Young, Large | 0.04 (-0.06, 0.14) | 0.412 |
| Old, Small | -0.05 (-0.10, -0.00) | 0.039 |
| Old, Large | -0.11 (-0.16, -0.06) | <0.001 |
| **Age-Size Group Comparison** | **Difference in Age Effect (95% CI)** | **Comparison p-value** |
| Old vs Young | -0.12 (-0.32, 0.06) | 0.189 |
| Old Dogs: Large vs Small | -0.06 (-0.12, -0.00) | 0.042 |
| Young Dogs: Large vs Small | 0.12 (-0.06, 0.29) | 0.196 |
